# Predicting Gastrointestinal Absorption of Prodrugs and their Drugs with the ANDROMEDA by Prosilico Software

**DOI:** 10.1101/2022.11.23.517725

**Authors:** Urban Fagerholm, Sven Hellberg, Jonathan Alvarsson, Ola Spjuth

**Affiliations:** Prosilico AB, Lännavägen 7, SE-141 45 Huddinge, Sweden; Department of Pharmaceutical Biosciences and Science for Life Laboratory, Uppsala University, Box 591, SE-751 24 Uppsala, Sweden

## Abstract

**Introduction:** Some prodrugs are developed in order to improve gastrointestinal absorption properties such as permeability and solubility/dissolution. Prediction of the uptake of prodrugs and their drugs is challening for reasons including gastrointestinal hydrolysis and active transport.

**Objective and Methodology:** The objective was to use the ANDROMEDA software by Prosilico to predict absorption characteristics - passive fraction absorbed (f_a,passive_), dose-adjusted dissolution potential (f_diss_) and total f_a_ (f_a_) - of prodrugs and their drugs (including drugs and their active metabolites), and to evaluate how they differ between prodrugs and drugs and the predictive accuracy of the software.

**Results:** 70 prodrug-drug pairs were found and selected for the study. The mean predicted f_a,passive_ and f_diss_ for the prodrugs were 0.74 and 0.94, respectively. Corresponding estimates for the drugs were 0.72 and 0.98, respectively. For non-hydrolyzed prodrugs, the median relative and absolute prediction errors for f_a_ were 1.17-fold and 0.08, respectively. Corresponding values for drugs were 1.11-fold and 0.07, respectively. The correlation between predicted and observed f_a_ for non-hydrolyzed ester prodrugs and drugs combined (predictive accuracy) was 0.6.

**Conclusion:** Prodrugs and drugs had similar average predicted f_a,passive_ and f_diss_, and most had or were predicted to have at least 50 % f_a_. The f_a_ for about 1/3 of non-hydrolyzed prodrugs was higher than for corresponding drugs, showing successful prodrug design. Adequate prediction accuracy validates ANDROMEDA for prediction of prodrug and drug absorption in man.

## Introduction

Prodrugs are developed in order to decrease local side effects and prolong patents of drugs and to improve their pharmacokinetic (PK) properties, for example, enhancing gastrointestinal absorption characteristics, such as permeability (P_e_) and solubility/dissolution (and thereby, increasing the oral bioavailability), and prolonging the elimination half-life (Fagerholm and Björnsson 2005; Rautio et al. 2018).

It has been approximated that 10 % of all marketed drugs can be considered prodrugs (Rautio et al. 2018). Many of these are esters that undergo hydrolysis in the gastrointestinal tract, which makes predictions of human *in vivo* gastrointestinal uptake challenging/difficult. The absorption of prodrugs designed to utilize intestinal uptake transporters is also challenging to predict.

Prediction of the gastrointestinal uptake requires prediction of absorption parameters/determinants such as passive Pe-driven fraction absorbed (f_a,passive_), active efflux by transporters such as MDR-1, BCRP and MRP2, active influx by tranporters such as PEPT1 (utilized by cephalosporins, penicillins, ACE-inhibitors, protease inhibitors, valacyclovir and levodopa) and *in vivo* dissolution potential (f_diss_; the maximum fraction dissolved).

The ANDROMEDA software by Prosilico for prediction, simulation and optimization of human clinical PK (with a major domain for compounds with molecular weight (MW) from 100 to 700 g/mole) has been sucessfully applied and validated for prediction of absorption/biopharmaceutics of small compounds in man in many studies (Fagerholm et al. 2022a–g). The software predicts f_a,passive_ (70 % predictions within ca ±7 %), f_diss_ (70 % predictions within ca ±12 %), efflux by MDR-1, BCRP and MRP2, and f_a_.

The objective of this study was to use ANDROMEDA to predict absorption characteristics (f_a,passive_, dose-adjusted f_diss_ and f_a_) of prodrugs and their drugs (including drugs and their active metabolites), and to evaluate how they differ between prodrugs and drugs and the predictive accuracy of the software.

## Methodology

ANDROMEDA by Prosilico was used to predict f_a,passive_, dose-adjusted f_diss_ (50 mg oral dose is the default for predicted f_diss_) and f_a_ for selected prodrugs and their drugs (or drugs and their active metabolites). Prodrug-drug pairs were searched for in the literature and the aim was to find at least 50 pairs of various kinds of prodrugs (including hydrolysis sensitive esters that are degraded to some extent before gastrointestinal uptake). No prodrug was actively excluded during this search process. Degradation in gastrointestinal fluids were not considered in the predictions. Observed/measured human *in vivo* f_a_-data were taken from various sources (Pham-The et al. 2013; Varma et al. 2010; Skolnik et al. 2010; Lin et al. 2011; Thomas et al. 2005; Matsson et al. 2005, Sköld et al. 2006; and data proprietary of Prosilico). Normally, compounds and data from training sets are not included in validation of the software. In this study, however, training set compounds and data were allowed in order to get better estimations of differences in absorption for each prodrug-drug (or drug-active metabolite) pair.

## Results & Discussion

70 prodrug-drug pairs (including many ester prodrugs and their drugs) were found and selected for the study (Table 1). Their MWs ranged from 130 to 744 g/mole. Data/compounds from training sets of prediction models were available in 32 cases (out of 140 cases), which is anticipated to have influenced the overall results to some minor extent.

**Table 1.** The 70 prodrug-drug (alternatively drug-active metabolite) pairs selected for the study.

| **Prodrug** | **Drug** |
| --- | --- |
| 6-monoacetylmorphine | Morphine |
| Abiratarone acetate | Abiratarone |
| Adefovir dipivoxil | Adefovir |
| Amitriptyline | Nortriptyline |
| Ampiroxicam | Piroxicam |
| Aripiprazole lauroxil | Aripiprazole |
| Azathioprine | Mercaptopurine |
| AZD3582 (naproxenol) | Naproxen |
| Azilsartan medoximil | Azilsartan |
| Bambuterol | Terbutaline |
| Becampicillin | Ampicillin |
| Caffeine | Theophylline |
| Candesartan cilexetil | Candesartan |
| Capecitabine | 5-FU |
| Carisoprodol | Meprobamate |
| Cefditoren pivoxil | Cefditoren |
| Cefotiam hexetil | Cefotiam |
| Cefpodoxime proxetil | Cefpodoxime |
| Ceftaroline fosamil | Caftaroline |
| Cefuroxime axetil | Cefuroxime |
| Chloramphenicol succinate | Chloramphenicol |
| Codeine | Morphine |
| Cortisone | Hydrocortisone |
| Cyclophosphamide | Aldophosphamide |
| Dabigatran etexilate | Dabigatran |
| Diazepam | Oxazepam |
| Enalapril | Enalaprilat |
| Eslicarbazepine acetate | Eslicarbazepine |
| Estramustine phosphate | Estramustine |
| Famciclovir | Penciclovir |
| Fenofibrate | Fenofibric acid |
| Fludarabine phosphate | Fludarabine |
| Fosamprenavir | Amprenavir |
| Fosaprepitant | Aprepitant |
| Fosfluconazole | Fluconazole |
| Fosphenytoin | Phenytoin |
| Fospropofol | Propofol |
| Gabapentin enacarbil | Gabapentin |
| Haloperidol decanoate | Haloperidol |
| Heroin | Morphine |
| Hetacillin | Ampicillin |
| Metoxibutropate | Ibuprofen |
| Imipramine | Desipramine |
| Irinotecan | SN-38 |
| Lenampicillin | Ampicillin |
| Levodopa | Dopamine |
| Lisdexamphetamine | Dextroamphetamine |
| Loratadine | Desloratadine |
| Midodrine | Desglymidodrine |
| Mycophenolate mofetil | Mycophenolic acid |
| Olmesartan medoxomil | Olmesartan |
| Paliperidone | Risperidone |
| Parecoxib | Valdecoxib |
| Pivampicillin | Ampicillin |
| Prednisone | Prednisolone |
| Primidone | Phenobarbital |
| Psilocybin | Psilocin |
| Rolitetracycline | Tetracycline |
| Salicylic acid | Acetylsalicylic acid |
| Serdexmethylphenidate | Dexmethylphenidate |
| Sulfasalazine | 5-aminosalicylic acid |
| Sulindac | Sulindac sulfide |
| Talampicillin | Ampicillin |
| Tedizolid phosphate | Tedizolid |
| Tenofovir disoproxil | Tenofovir |
| Terfenadine | Fexofenadine |
| Valaciclovir | Aciclovir |
| Valganciclovir | Ganciclovir |
| Valpromide | Valproic acid |
| Ximelagatran | Melagatran |

The mean predicted f_a,passive_ and f_diss_ for the prodrugs were 0.74 (range 0.02-1.00) and 0.94 (range 0.32-1.00), respectively (Figures 1 and 2). Corresponding estimates for the drugs were 0.72 (range 0.02-1.00) and 0.98 (range 0.65-1.00), respectively. 29 (41 %) and 7 (10 %) prodrugs had predicted f_a,passive_≥0.9 and <0.25, respectively. For drugs, corresponding values were 31 (44 %) and 7 (10 %), respectively. 29 (41 %) and 7 (10 %) prodrugs had predicted f_a,passive_≥0.9 and <0.25, respectively. For drugs, corresponding values were 31 (44 %) and 7 (10 %), respectively. 59 (84 %) and 6 (9 %) prodrugs had predicted f_diss_≥0.9 and <0.7, respectively. For drugs, corresponding values were 66 (94 %) and 1 (1 %), respectively.

**Figure 1.**
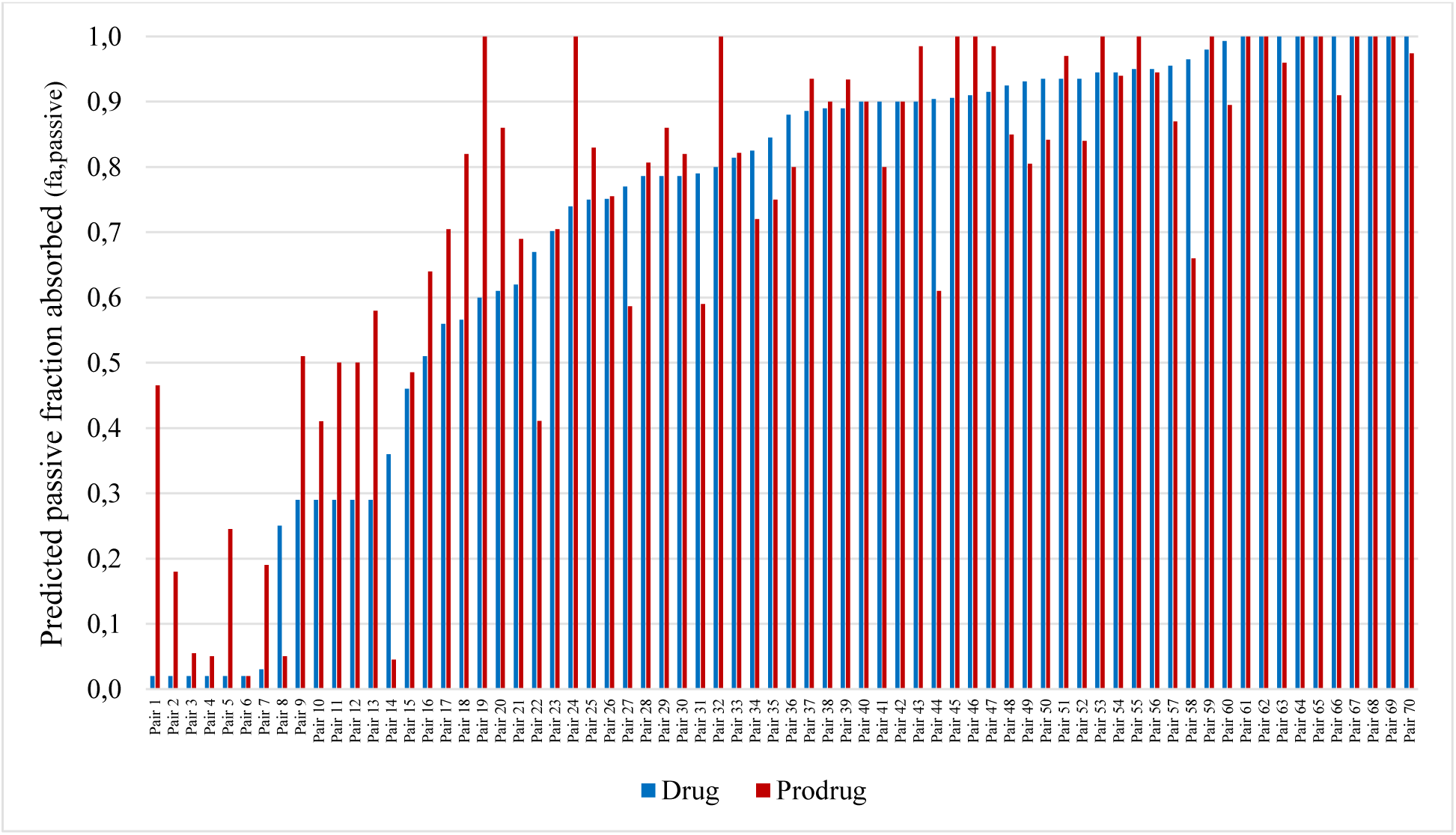
Ranked predicted passive fraction absorbed (f_a,passive_) for 70 prodrug-drug pairs. Ranking was based on data for drugs.

**Figure 2.**
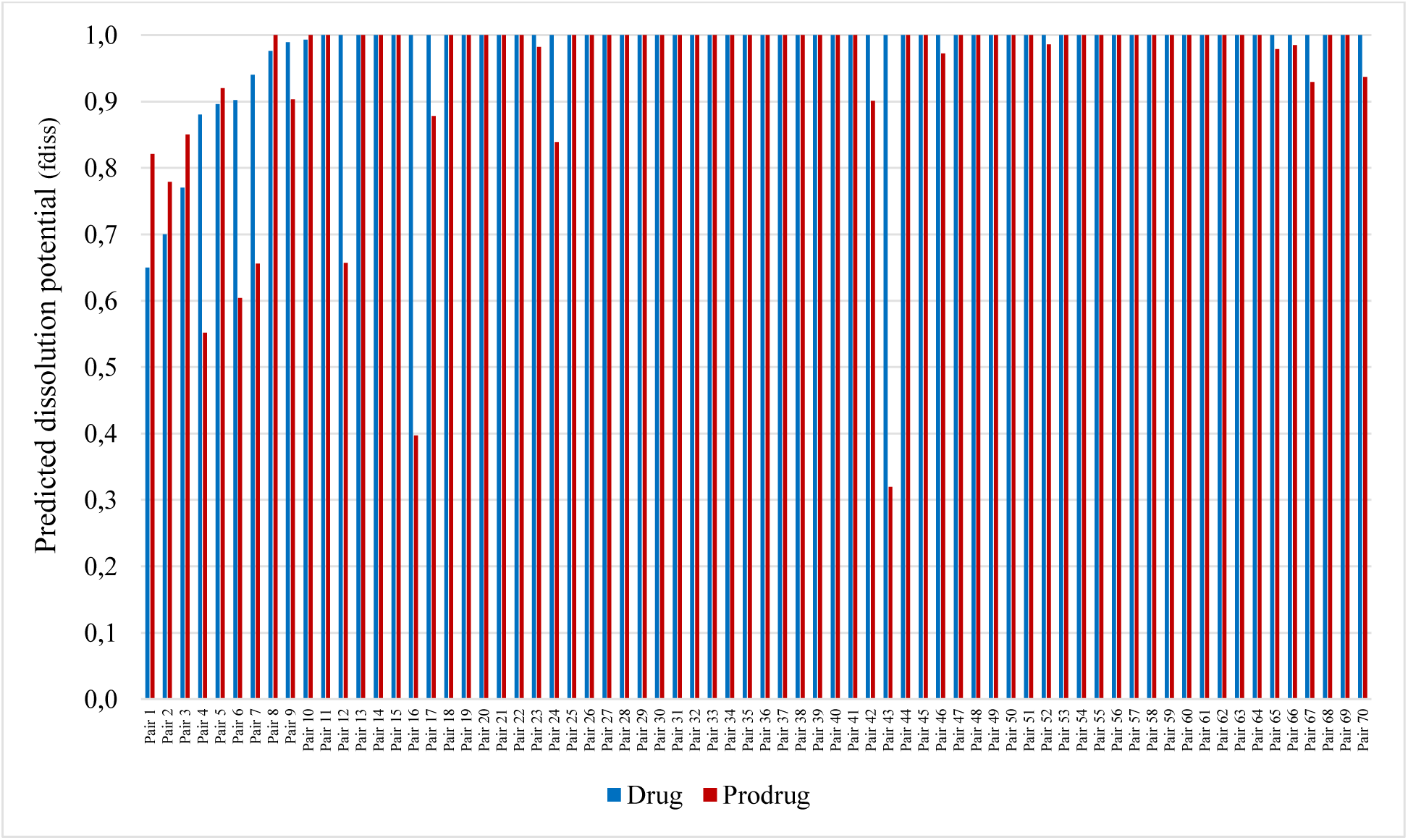
Ranked predicted maximum dissolution potential (f_diss_) for 70 prodrug-drug pairs. Ranking was based on data for drugs.

Thus, the majority of the selected prodrugs and drugs have good/adequate predicted passive P_e_ and f_diss_.

For prodrugs, the median (mean) relative and absolute prediction errors for f_a_ were 1.17 (1.24)-fold and 0.08 (0.15) (n=17; non-hydrolyzed prodrugs with observed estimates only), respectively (Table 2). For 7 prodrugs with approximated f_a_ (for example, moderate, >0.90 and >0.26-0.91), the median relative maximum prediction error for f_a_ was 1.69-fold. Only 1 to 4 (4 to 16 %) of the prodrugs had or were predicted to have a f_a_ of less than 50 %.

**Table 2.** *In silico* prediction results for 25 non-hydrolyzed prodrugs.

| **Prodrug** | **Observed**<br>**$f_a$** | **Predicted**<br>**$f_a$** | **Error** | **Max**<br>**error** | **Abs**<br>**error** |
| --- | --- | --- | --- | --- | --- |
| Adefovir dipivoxil | >0,6 | 1,00* (0,47) |  | 1,67 |  |
| Amitriptyline | 0,95 | 1,00 | 1,05 |  | 0,05 |
| Azathioprine | 0,88 | 0,46 | 1,90 |  | 0,42 |
| Caffeine | 1,00 | 0,76 | 1,32 |  | 0,24 |
| Capecitabine | Mod/high | 0,59 |  | 1,69 |  |
| Cefpodoxime proxetil | Poor | 0,03 |  |  |  |
| Codeine | 0,93 | 0,69 | 1,36 |  | 0,24 |
| Cortisone | >0,26-0,91 | 0,94 |  | 3,48 |  |
| Cyclophosphamide | 0,90 | 0,76 | 1,19 |  | 0,15 |
| Diazepam | 1,00 | 1,00 | 1,01 |  | 0,01 |
| Enalapril | 0,65 | 0,92* (0,15) | 1,38 |  |  |
| Fenofibrate | >0,90 | 0,99 |  | 1,09 |  |
| Imipramine | 1,00 | 1,00 | 1,00 |  | 0,00 |
| Irinotecan | >0,09 | 0,46 |  | 4,60 |  |
| Levodopa** | 1,00 | 1,00* (0,41) | 1,00 |  |  |
| Loratadine | 0,90 | 0,92 | 1,02 |  | 0,02 |
| Midodrine | 0,93 | 0,51 | 1,82 |  | 0,42 |
| Paliperidone | >0,28 | 0,68 |  | 2,34 |  |
| Prednisone | 0,95 | 0,93 | 1,02 |  | 0,02 |
| Primidone | 0,75 | 0,97 | 1,30 |  | 0,22 |
| Psilocybin | >0,53 | 0,65 |  | 1,54 |  |
| Sulindac | 0,90 | 0,84 | 1,07 |  | 0,06 |
| Terfenadine | 1,00 | 0,95 | 1,06 |  | 0,05 |
| Valaciclovir | 0,68 | 0,92* (0,64) | 1,35 |  | 0,24 |
| Ximelagatran | 0,55 | 0,47 | 1,17 |  | 0,08 |
| Min |  |  | 1,00 | 1,09 | 0,00 |
| Max |  |  | 1,90 | 4,60 | 0,42 |
| Mean |  |  | 1,24 | 2,34 | 0,15 |
| Median |  |  | 1,17 | 1,69 | 0,08 |

\*Considering active uptake (passive contribution within parentheses)
\*\*Stabilized with the dopa decarboxylase inhibitor benserazide

For drugs, the median (mean) relative and absolute prediction errors for f_a_ were 1.11 (1.63)- fold and 0.07 (0.11) (n=45; drugs (or active metabolites) with observed estimates only), respectively (Table 3). For 12 drugs with approximated f_a_ the median relative maximum prediction error for f_a_ was 1.49-fold. 6 to 8 (13 or 18 %) of the drugs had or were predicted to have f_a_<50 %.

**Table 3.** *In silico* prediction results for 45 drugs.

| Drug | Observed<br>$f_a$ | Predicted<br>$f_a$ | Error | Max error | Abs error |
| --- | --- | --- | --- | --- | --- |
| 5-aminosalicylic acid | High | 0,61 |  | 1,64 |  |
| 5-FU | High | 0,57 |  | 1,76 |  |
| Acetylsalicylic acid | 0,82 | 0,76 | 1,08 |  | 0,06 |
| Aciclovir | 0,23 | 0,41 | 1,80 |  | 0,18 |
| Adefovir | 0,16 | 0,21* (0,01) | 1,31 |  | 0,05 |
| Ampicillin | 0,65 | 0,91* (0,18) | 1,40 |  | 0,26 |
| Amprenavir | 0,70 | 0,87 | 1,24 |  | 0,17 |
| Aprepitant | >0,60-0,65 | 0,58 |  | 1,72 |  |
| Aripiprazole | >0,87 | 0,88 |  | 1,13 |  |
| Cefuroxime | 0,44 | 0,37* (0,02) | 1,19 |  | 0,07 |
| Chloramphenicol | 0,89 | 0,59 | 1,52 |  | 0,30 |
| Desglymidodrine | >0,5 | 0,64 |  | 1,56 |  |
| Desipramine | 1,00 | 1,00 | 1,00 |  | 0,00 |
| Dexmethylphenidate | 0,90 | 0,90 | 1,00 |  | 0,00 |
| Dextroamphetamine | >0,75 | 0,93 |  | 1,07 |  |
| Enalaprilat | 0,15 | 0,01 | 12,50 |  | 0,14 |
| Estramustine | 0,83 | 0,61 | 1,36 |  | 0,22 |
| Fenofibric acid | >0,81 | 0,78 |  | 1,28 |  |
| Fexofenadine | 0,58 | 0,76 | 1,31 |  | 0,18 |
| Fluconazole | 0,95 | 0,67 | 1,40 |  | 0,27 |
| Gabapentin | >0,65 | 0,70 |  | 1,42 |  |
| Ganciclovir | >0,06 | 0,30 |  | 3,29 |  |
| Haloperidol | 0,95 | 1,00 | 1,05 |  | 0,05 |
| Hydrocortisone | 0,91 | 0,89 | 1,03 |  | 0,02 |
| Ibuprofen | 0,97 | 0,91 | 1,07 |  | 0,06 |
| Melagatran | 0,15 | 0,37 | 2,48 |  | 0,22 |
| Mercaptopurine | 0,50 | 0,50 | 1,01 |  | 0,01 |
| Morphine | 0,85 | 0,79 | 1,08 |  | 0,06 |
| Mycophenolic acid | >0,4 | 0,39 |  | 2,59 |  |
| Naproxen | 0,99 | 0,85 | 1,17 |  | 0,14 |
| Nortriptyline | 1,00 | 1,00 | 1,00 |  | 0,00 |
| Olmesartan | >0,26 | 0,82 |  | 1,22 |  |
| Oxazepam | 0,95 | 0,94 | 1,02 |  | 0,01 |
| Penciclovir | 0,10 | 0,34 | 3,43 |  | 0,24 |
| Phenobarbital | 1,00 | 1,00 | 1,00 |  | 0,00 |
| Phenytoin | 0,90 | 1,00 | 1,11 |  | 0,10 |
| Piroxicam | 1,00 | 1,00 | 1,00 |  | 0,00 |
| Prednisolone | 0,99 | 0,89 | 1,11 |  | 0,10 |
| Propofol | 1,00 | 0,99 | 1,01 |  | 0,01 |
| Psilocin | >0,5 | 0,82 |  | 1,22 |  |
| Risperidone | 0,99 | 0,80 | 1,23 |  | 0,19 |
| Terbutaline | 0,73 | 0,57 | 1,29 |  | 0,16 |
| Tetracycline | 0,83 | 0,90* (0,42) | 1,08 |  | 0,07 |
| Theophylline | 1,00 | 0,76 | 1,32 |  | 0,24 |
| Valproic acid | 0,99 | 0,95 | 1,04 |  | 0,04 |
| Min |  |  | 1,00 | 1,07 | 0,00 |
| Max |  |  | 12,50 | 3,29 | 0,30 |
| Mean |  |  | 1,63 | 1,66 | 0,11 |
| Median |  |  | 1,11 | 1,49 | 0,07 |

\*Considering active uptake (passive contribution within parentheses)

The correlation (Q^2^; predictive accuracy) between predicted and observed f_a_ was 0.62 (observed f_a_ = 0.84 • predicted f_a_ + 0.15; n=50; data for both non-hydrolyzed ester prodrugs and drugs combined; Figure 3). Removal of the active uptake component for 7 compounds resulted in a higher intercept (0.33), but similar Q^2^ (0.57).

**Figure 3.**
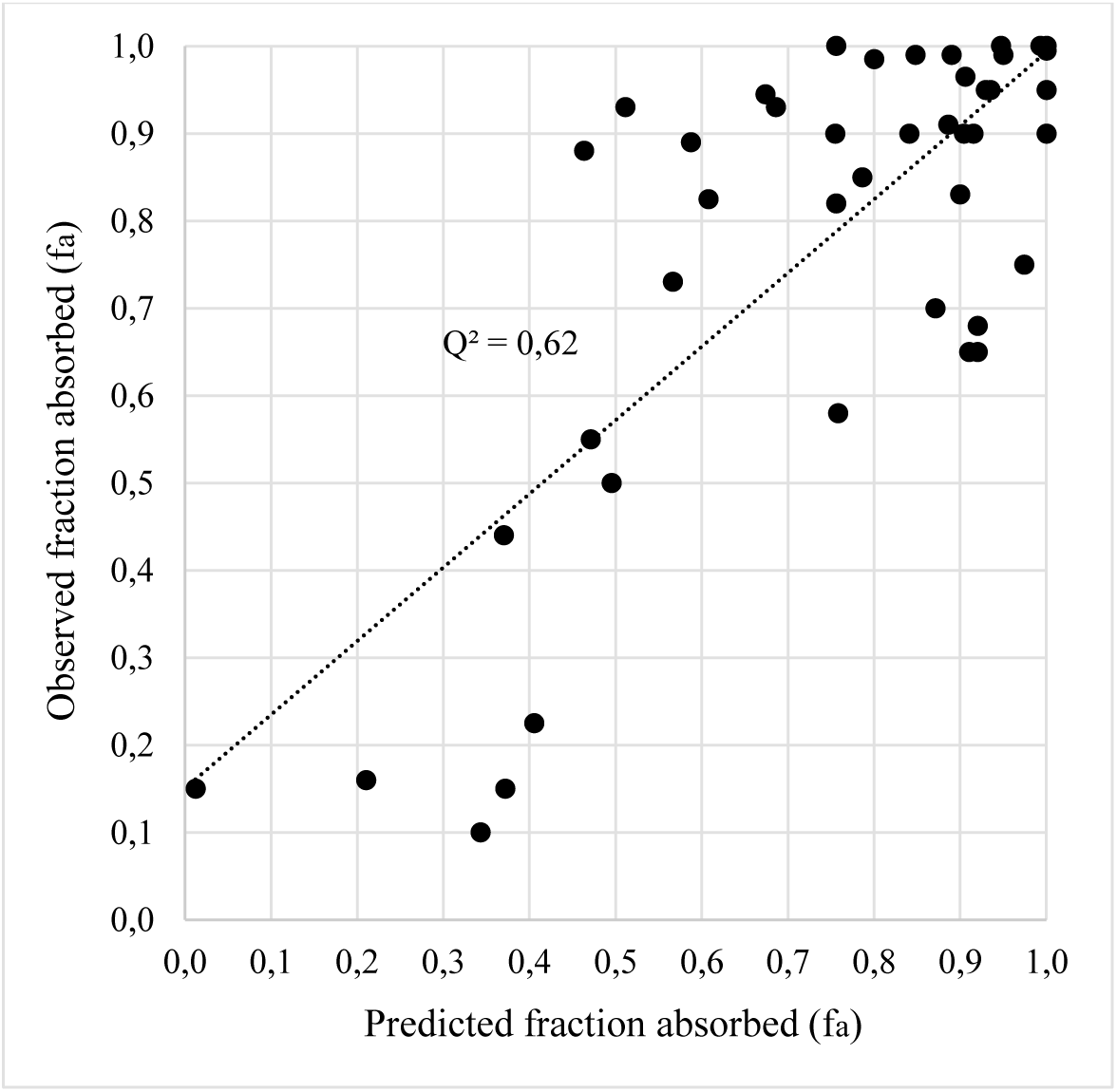
*In silico* predicted *vs* observed fraction absorbed (fa) for 50 prodrugs and drugs.

Among 25 pairs with observed and/or predicted f_a_ for both the prodrug (excluding hydrolyzed esters) and the drug, 8 (32 %) showed apparently higher f_a_ for the prodrug (Table 4). These were adefovir dipivoxil (active uptake), enalapril (active uptake; bioavailability of enalaprilat increased from 3-12% to 60-70% on dosing enalapril), fenofibrate (higher f_a,passive_ and possibly removed efflux), levodopa (active uptake), loratadine (higher f_a,passive_ and f_diss_), terfenadine (higher f_a,passive_), valaciclovir (active uptake; 3-5 times higher oral bioavailability of acyclovir following valaciclovir dosing) and ximelagatran (higher f_a,passive_).

**Table 4.** *In silico* prediction results for 25 prodrug-drug pairs (excluding hydrolyzed esters).

| **Prodrug** | **Observed**<br>**fa** | **Predicted**<br>**fa** | **Drug** | **Observed**<br>**fa** | **Predicted**<br>**fa** | **Prodrug**<br>**>Drug** |
| --- | --- | --- | --- | --- | --- | --- |
| Adefovir dipivoxil | >0,6 | 1,00* (0,47) | Adefovir | 0,16 | 0,21* (0,01) | Yes |
| Amitriptyline | 0,95 | 1,00 | Nortriptyline | 1,00 | 1,00 |  |
| Azathioprine | 0,88 | 0,46 | Mercaptopurine | 0,50 | 0,50 |  |
| Caffeine | 1,00 | 0,76 | Theophylline | 1,00 | 0,76 |  |
| Capecitabine | Mod/high | 0,59 | 5-FU | High | 0,57 |  |
| Cefpodoxime proxetil | Poor | 0,03 | Cefpodoxime | - | 0,01 |  |
| Codeine | 0,93 | 0,69 | Morphine | 0,85 | 0,79 |  |
| Cortisone | >0,26-0,91 | 0,94 | Hydrocortisone | 0,91 | 0,89 |  |
| Cyclophosphamide | 0,90 | 0,76 | Aldophosphamide | - | 0,75 |  |
| Diazepam | 1,00 | 1,00 | Oxazepam | 0,95 | 0,94 |  |
| Enalapril | 0,65 | 0,92* (0,15) | Enalaprilat | 0,15 | 0,01 | Yes |
| Fenofibrate | >0,90 | 0,99 | Fenofibric acid | >0,81 | 0,78 | Yes |
| Imipramine | 1,00 | 1,00 | Desipramine | 1,00 | 1,00 |  |
| Irinotecan | >0,09 | 0,46 | SN-38 | - | 0,66 |  |
| Levodopa** | 1,00 | 1,00* (0,41) | Dopamine | - | 0,47 | Yes |
| Loratadine | 0,90 | 0,92 | Desloratadine | - | 0,69 | Yes |
| Midodrine | 0,93 | 0,51 | Desglymidodrine | >0,5 | 0,64 |  |
| Paliperidone | >0,28 | 0,68 | Risperidone | 0,99 | 0,80 |  |
| Prednisone | 0,95 | 0,93 | Prednisolone | 0,99 | 0,89 |  |
| Primidone | 0,75 | 0,97 | Phenobarbital | 1,00 | 1,00 |  |
| Psilocybin | >0,53 | 0,65 | Psilocin | >0,5 | 0,82 |  |
| Sulindac | 0,90 | 0,84 | Sulindac sulfide | - | 0,83 |  |
| Terfenadine | 1,00 | 0,95 | Fexofenadine | 0,58 | 0,76 | Yes |
| Valaciclovir | 0,68 | 0,92* (0,64) | Aciclovir | 0,23 | 0,41 | Yes |
| Ximelagatran | 0,55 | 0,47 | Melagatran | 0,15 | 0,37 | Yes |

\*Considering active uptake (passive contribution within parentheses)
\*\*Stabilized with the dopa decarboxylase inhibitor benserazide

Limitations with the study include the fact that all available prodrugs were not selected and uncertainties regarding active uptake and gastrointestinal hydrolysis. Inclusion of compensation for active uptake was done in retrospect (when active transport was known). There are also uncertainties/variability of collected *in vivo* f_a_-data. Despite this, prediction results are encouraging.

In conclusion, prodrugs did not have larger or smaller average predicted f_a,passive_ and f_diss_ than their drugs. However, the f_a_ for about 1/3 of non-hydrolyzed prodrugs was higher than for corresponding drugs, showing successful prodrug design. Most prodrugs and drugs had or were predicted to have at least 50 % f_a_. The overall prediction accuracy was adequate, which validates ANDROMEDA for prediction of prodrug and drug absorption in man.

